# Challenging the function of motor inhibition: Does it really assist action selection?

**DOI:** 10.1101/504159

**Authors:** Caroline Quoilin, Fanny Fievez, Julie Duque

**Affiliations:** Institute of Neuroscience, Université catholique de Louvain, Brussels, Belgium.; Laboratory for Experimental Psychopathology, Psychological Sciences Research Institute, Université catholique de Louvain, Louvain-la-Neuve, Belgium.

**Keywords:** Inhibitory control, Transcranial Magnetic Stimulation (TMS), Action preparation, Preparatory inhibition, Motor-evoked potential (MEP)

## Abstract

By applying transcranial magnetic stimulation (TMS) over the primary motor cortex (M1) to elicit motor-evoked potentials (MEPs) in muscles of the contralateral hand during reaction time (RT) tasks, many studies have reported a strong suppression of MEPs during action preparation, a phenomenon called preparatory inhibition. Several hypotheses have been put forward regarding the role of this inhibition, with the predominant view suggesting that it would help action selection. However, this assumption is still a matter of debate. Here, we aimed at directly addressing this idea by comparing MEPs in a task that required subjects to select a finger response within a set of predefined options (choice RT task: left or right index finger abduction) or when subjects simply had to provide the same finger response on every trial, in the absence of choice (simple RT task). Moreover, we minimized any effect that could be associated with other forms of inhibition. In both versions of the task, TMS was applied on both M1 (double-coil protocol) at several time points between the go signal and the left or right index finger response, eliciting MEPs bilaterally in the prime mover (index finger agonist) and in an irrelevant muscle (pinky agonist). Overall, MEP suppression was moderate in this study; it was only found for the irrelevant muscle. As such, MEPs in the index agonist were facilitated when elicited in a responding hand (e.g. left MEPs preceding left responses) and remained mostly unchanged in a non-responding hand (e.g. left MEPs preceding right responses). In contrast, MEPs were almost always suppressed in the pinky muscle when elicited in the non-responding hand and sometimes also in the responding hand. Importantly, this effect was more consistent in the choice than in the simple RT task, supporting the view that preparatory inhibition may assist action selection. Moreover, the fact that it sometimes concerned the responding hand is coherent with the idea of a global process, suppressing broadly the motor system.

**Highlights:**

- Addressed the debated idea that preparatory inhibition assists action selection
- Compared MEPs in simple and choice RT tasks (no catch, no delay)
- Detected preparatory suppression of MEPs, although weaker than usual
- Observed a more consistent suppression of MEPs in the choice than simple setting
- Provided support for action selection hypothesis, with traces of broad inhibition

## 1. Introduction

Action preparation relies on the operation of control processes that leave their imprint on the motor system (Alamia et al., 2018; Derosiere et al., 2018; Lebon et al., 2018; Thura and Cisek, 2014). Interestingly, these modulatory changes can be examined in humans by applying transcranial magnetic stimulation (TMS) over the primary motor cortex (M1). When applied over M1, single-pulse TMS elicits motor-evoked potentials (MEPs) in targeted contralateral muscles, and the amplitude of these MEPs reflects the excitability of the corticospinal pathway at the time of stimulation (Bestmann and Duque, 2016; Hannhah and Rothwell, 2017). Hence, by applying TMS at different moments preceding movement onset, one can track changes in corticospinal excitability (CSE) occurring along with action preparation. Using this approach, a considerable amount of studies has reported a strong suppression of MEPs (reviewed in Bestmann and Duque, 2016). That is, in many circumstances, MEP amplitudes are found to be smaller when an action is being prepared compared to when MEPs are elicited in a baseline condition, a phenomenon called preparatory inhibition (Duque et al., 2017). Several hypothesis have been put forward regarding the role of motor suppression during action preparation (Duque et al., 2010; Greenhouse et al., 2015; Hannah et al., 2018; Kaufman et al., 2014). Among them, a predominant view is that preparatory inhibition serves to assist action selection, consistent with a contribution of inhibition to the generation of goal-oriented behaviors (Bestmann and Duque, 2016).

The idea that preparatory inhibition helps action selection was motivated, in part, by early TMS studies showing consistent MEP suppression in non-selected effectors during choice reaction time (RT) tasks (Duque et al., 2005; Leocani et al., 2000). In such tasks, subjects are usually required to select a response within a set of predefined options and to release it as fast as possible following an imperative signal. The choice is often between a finger movement of the left or the right hand and TMS is usually applied during the RT period, eliciting MEPs in the targeted finger muscle at various times preceding movement onset (Duque et al., 2014; Klein et al., 2016). Depending on whether subjects prepare a left or a right hand response, these MEPs will either fall in a (selected) responding hand (e.g. Left MEPs elicited in left hand trials) or in a (non-selected) non-responding hand (e.g. Left MEPs elicited in right hand trials). MEP amplitudes typically increase in the former condition, while, as mentioned above, they are often suppressed in the non-responding hand (Burle et al., 2004; Duque et al., 2005; Leocani et al., 2000).

The action selection hypothesis is supported by the fact that the strength of this MEP suppression increases with the risk of selecting an inappropriate response. This may arise because of incongruent sensory information (Burle et al., 2016; Duque et al., 2016; Klein et al., 2014) or because a non-selected response is prepotent (Klein et al., 2016; Meckler et al., 2011). For instance, in right-handers, MEPs elicited close to movement onset are more suppressed in the right than in the left non-responding hand, possibly allowing to prevent inadvertent movements with the right dominant hand following cues requiring left hand responses (Klein et al., 2016). Furthermore, the amount of MEP suppression depends on the anatomical and/or functional relationship between the competing effectors (Duque et al., 2014; Labruna et al., 2014), further indicating a link between preparatory inhibition and the selection aspects of action preparation.

However, other recent findings cast doubt on such role of motor inhibition in action selection. First, a large body of research has shown that, in many variants of the choice RT task, MEP suppression is already present before the imperative signal, especially when a fixation cross announces the beginning of the trial or when a preparatory cue indicates (part of) the required movement in advance (reviewed in Duque et al., 2017). Importantly, this pre-imperative inhibition is rather global, concerning all types of effectors, whether they are (potentially) selected, non-selected or even irrelevant in the task (Derosiere, 2018; Duque et al., 2014; Greenhouse et al., 2015; Quoilin et al., 2016). More critically, at that time, the MEP suppression is substantial even when subjects have to perform the same response on every trial in simple RT tasks (Greenhouse et al., 2015). Hence, pre-imperative inhibition also occurs in the absence of selection requirements and may persist over time, potentially accounting for the MEP suppression observed in non-responding effectors at later time points preceding movement onset. Clearly, this idea contrasts with the view that preparatory inhibition would be deployed during action selection to solve a competition between potential effectors. Finally, Greenhouse et al. (2015) observed inhibition in the nonresponding effector during a simple RT task even when the imperative signal was not preceded by a preparatory cue. However, in that study, the authors included a large amount of catch trials, in which the imperative signal was not presented (Quoilin and Derosiere, 2015). This aspect of the design may have increased proactive control, which is known to exert global inhibitory effects on the motor system (Greenhouse et al., 2012; Wessel et al., 2016).

Hence, at this point, whether aspects of preparatory inhibition contribute to action selection is still unclear. Here, we aimed at addressing this question by directly comparing MEP changes in simple and choice RT tasks, while minimizing any effect that could be associated with other forms of inhibition. Participants were required to respond with the left or the right index finger as fast as possible following the onset of an imperative signal. In the simple RT blocks, the finger response was fixed for the entire block, allowing to evaluate CSE in the absence of choice. By contrast, in the choice RT blocks, the appropriate response had to be selected on each trial, according to the imperative signal. Importantly, in order to reduce pre-imperative or proactive inhibition, the imperative signal was never preceded by a fixation cross or a preparatory cue, and the duration of the inter-trial interval was variable. Moreover, catch trials were not included. Based on the action selection hypothesis, we predicted that preparatory inhibition would be deeper during the postimperative period that precedes movement onset in the choice compared to the simple version of the RT task.

## 2. Material and Methods

### 2.1. Participants

A total of 18 right-handed volunteers (10 women; mean age = 22.4 ± 2.15 years old) participated in the experiment. Handedness was determined via a condensed version of the Edinburgh Handedness Inventory (Oldfield, 1971). None of the participants suffered from any neurological disorder or had a history of psychiatric illness, drug or alcohol abuse; none either was undergoing any drug treatment that could influence performance or neural activity. Participants were naive to the purpose of the study and were financially compensated. All gave written informed consent at the beginning of the study, in accordance with the protocol approved by the Ethics Committee of the Université catholique de Louvain (UCLouvain), in compliance with the principles of the Declaration of Helsinki.

### 2.2. The “Rolling Ball” Task

Participants sat in front of a computer screen, positioned about 60 cm in front of them, with both forearms resting in a semi-flexed position and the hands placed palms down on a response device. They performed a RT task, which was implemented with Matlab 7.5 (The Mathworks, Natick, Massachusetts, USA) and the Cogent 2000 toolbox (Functional Imaging Laboratory, Laboratory of Neurobiology and Institute of Cognitive Neuroscience at the Wellcome Department of Imaging Neuroscience, London, UK). The task consisted in a virtual “rolling ball” game, in which participants had to use their left or right index finger “to shoot a ball”, allowing it to roll over a bridge into a goal displayed on the screen (Grandjean et al., 2018; Quoilin et al., 2016, 2018; Vassiliadis et al., 2018; see Figure 1). The required response was indicated by the position of the ball: participants were instructed to respond with an abduction of the left index finger when the ball was presented on the left side of the screen, and with the right index finger when it appeared on the right.

**Figure 1.**
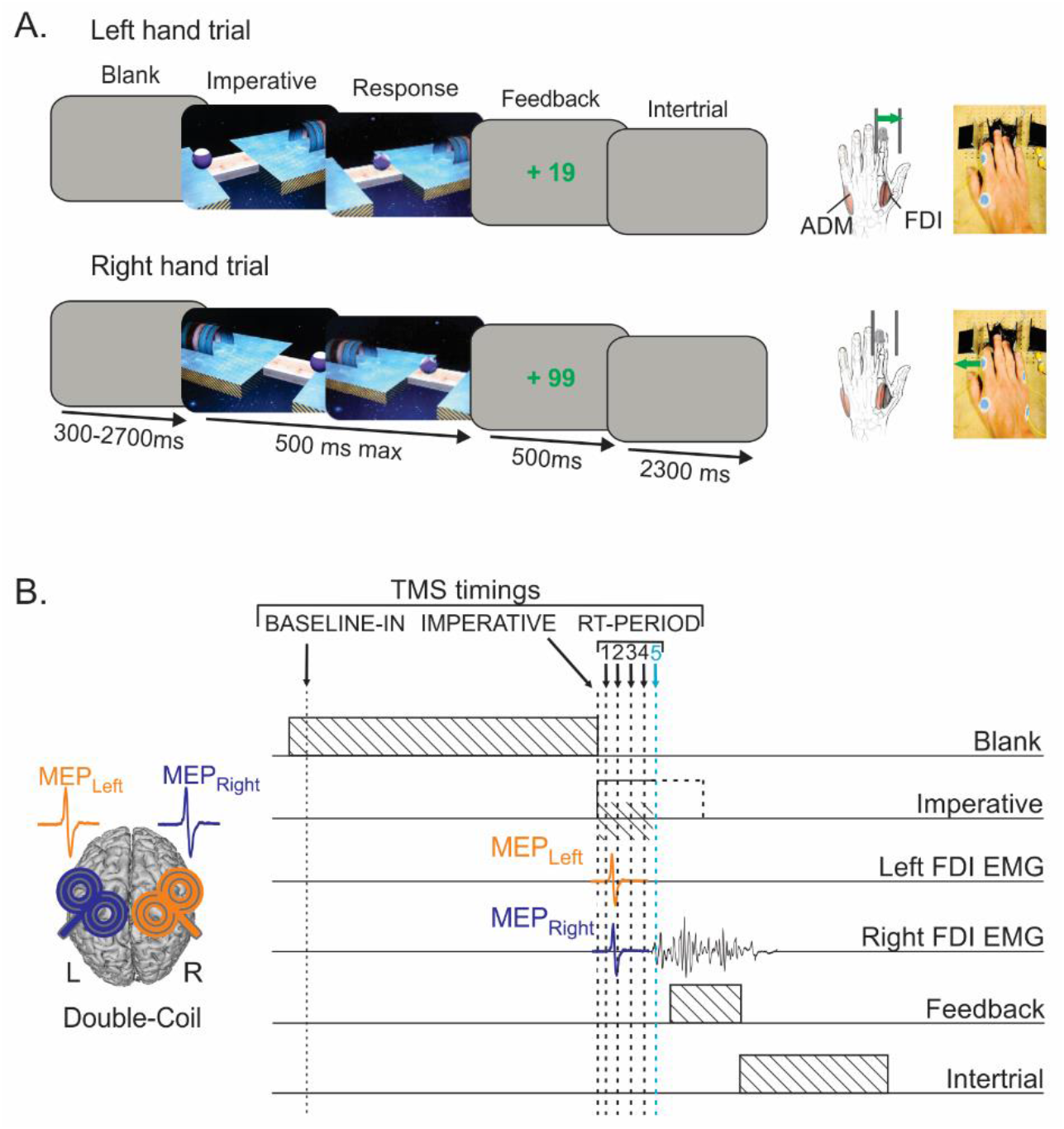
(A): Sequence of events in the simple and choice blocks of the reaction time (RT) task. Subjects were required to perform an abduction of the left or right index finger in order to “shoot a ball as fast as possible into a goal”. Each trial started with the presentation of a blank screen displayed for a random period of 300 to 2700 ms, followed by the onset of the imperative signal. This signal indicated the movement to perform, as participants had to respond with the left hand when the ball was displayed on the left side of the screen, and with the right hand when it was presented on the right. The imperative signal disappeared as soon as a finger response was detected (500 ms max) and a feedback score was then displayed for 500 ms. Finally, the inter-trial interval lasted for 2300 ms. Importantly, participants performed 4 simple RT blocks in which the ball always appeared on the same side of the screen (2 blocks for each side) and 4 choice RT blocks in which the ball could randomly appear on the left or the right side. (B): TMS protocol and timings. Motor-evoked potentials (MEPs) were elicited using a double-coil TMS method with pulses delivered to the left and right primary motor cortices at a near-simultaneous time (i.e. 1 ms interval). During each trial, TMS pulses were delivered at one out of six (simple blocks) or seven (choice blocks) possible timings. The pulses either occurred 200 ms after the blank screen onset (TMS_BASELINE-IN_), at the imperative signal onset (TMS_IMPERATIVE_), or during the RT period (TMS_RT-PERIOD1-5_, with the last timing only probed in the choice blocks). FDI = first dorsal interosseous. ADM = abductor digiti minimi.

Two variants of the rolling ball task were used in separate blocks of trials. That is, in some blocks, the ball systematically appeared on the same side of the screen. Hence, in this simple RT version of the task, no choice was required: subjects had to respond with the same index finger (left or right) in all trials of a given block. Besides, in other blocks, we used a choice RT version of the task where the ball appeared randomly on the left or on the right side of the screen, thus requiring subjects to select the appropriate finger response on each trial.

The sequence and timing of events are shown in Figure 1A (left side). Each trial started with the presentation of a blank screen (light grey) lasting for a random duration ranging from 300 to 2700 ms. Then, the imperative screen appeared, which consisted of the ball and the goal presented on separate platforms connected by a bridge. Importantly, the presentation of the bridge was not delayed with respect to the ball appearance (as in most previous studies on preparatory inhibition, see Quoilin et al., 2016, 2018; Vassiliadis et al., 2018), a situation that would have required subjects to postpone their finger response until the bridge onset. Rather, here participants could respond as soon as the ball appeared, by abducting the correct index finger within a maximum time of 500 ms. The imperative screen disappeared once a response was detected (or after 500 ms) and a feedback was presented for 500 ms. Following a correct response, the feedback consisted of a positive score depicted in green, which ranged from +5 to +100 and was inversely proportional to the trial’s RT 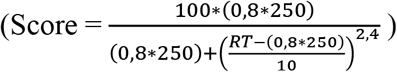. score of 5 occurred following slow responses provided in 500 ms (corresponding to the longest RT considered as acceptable) while a maximum score of 100 was obtained following fast responses provided in 200 ms or less. By contrast, participants were presented with a fixed negative score depicted in red when they responded with the incorrect finger (−20), when they provided their response too slowly (i.e. more than 500 ms after the onset of the imperative; −50), or when they responded prematurely (i.e. before the imperative signal onset; −75). Finally, the duration of the inter-trial interval was set at 2300 ms (blank screen). As a consequence, the time elapsed between the end of a trial and the onset of the imperative on the next trial was very variable, fluctuating between 2600 and 5000 ms. This variable delay, together with the absence of preparatory cue, made it hard for the subjects to predict the time of the imperative signal, reducing the risk of premature responses in the current study.

The participant responses were recorded using a device developed in our laboratory and designed to detect horizontal movements of the index fingers (Grandjean et al., 2018; Quoilin et al., 2016; Vassiliadis et al., 2018). It was composed of two pairs of metal edges fixed on wooden supports (one for each hand) and each response required to move one index finger from the outer to the inner metal edge (see Figure 1A, right side). The contact between the finger and the metal parts was continuously monitored using a Makey Makey printed circuit board with an ATMega32u4 microcontroller running the Arduino Leonardo firmware, based on the principle of high resistance switching between two electrical contacts. To enhance connectivity, a thimble was placed on both index fingers. This device provided us with a very precise measure of the RTs, defined as the time elapsed between the imperative signal onset and the moment at which the index finger left the outer metal edge (precision = 1 ms). Moreover, it allowed us to control for any anticipated movement. That is, the device permanently checked the initial position of each index finger (which had to be in contact with the outer metal edge) and any contact release before the onset of the imperative signal led to the cancellation of the trial and to a penalty (score of −75, see above).

### 2.3. Experimental procedure

The testing always began with a short practice period to allow participants to become familiar with the task. Then, the main phase of the experiment involved 8 blocks of 60 trials; half of which were run in the simple RT mode and the other half in the choice mode. For the simple RT blocks, the ball systematically appeared on the same side of the screen, always requiring either a left or a right index finger response on each trial (2 blocks for each hand). By contrast, the choice RT blocks consisted of an equal proportion of left and right hand trials, with the ball appearing randomly on the left or right side (4 blocks). The simple and choice RT conditions were tested in successive blocks but their order was counterbalanced between subjects. Blocks lasted approximatively 6-7 minutes, and participants were given a short break between each block. They were always informed about the type of block they would be performing next.

TMS elicited MEPs at different time points during the task (only one timing per trial; double-coil procedure, see below and Figure 1B). To establish a baseline measure of CSE in the simple and choice RT blocks, TMS was sometimes applied 200 ms after the onset of the initial blank screen (TMS_BASELINE-IN_; 10 MEPs/block). In other trials, the TMS pulse could also fall at the onset of the imperative signal (TMS_IMPERATIVE_; 10 MEPs/block); this timing was used to check for any change in CSE that could be due to the anticipation of the imperative signal. As mentioned above, we expected this effect to be marginal, especially in choice RT blocks, given the absence of preparatory cue and the variable time of the imperative signal onset. Moreover, several TMS pulses occurred during the RT period. In the simple RT blocks, these pulses were delivered 80, 130, 250 or 300 ms after the onset of the imperative signal (TMS_RT-PERIOD1-4_, 10 MEPs per time/block; total of 20 MEPs per time and responding side). In the choice RT blocks, we included an additional timing at 350 ms to account for the longer RTs in this condition (Greenhouse et al., 2015), in order to cover the whole RT period (TMS_RT-PERIOD1-5_; 4 MEPs per time and responding side/block; total of 16 MEPs per time and responding side). Finally, 20 TMS pulses were also applied between each block to obtain a baseline measure of CSE outside the context of the task (TMS_BASELINE-OUT_). Importantly, these TMS_BASELINE-OUT_ MEPs were grouped and averaged according to whether they surrounded a simple left, simple right or choice RT block.

### 2.4. TMS protocol

TMS was delivered using a double-coil method recently developed in our laboratory where both M1 are stimulated with a 1 ms inter-pulse interval (double-coil_1ms_ TMS), eliciting MEPs in both hands at a near simultaneous time (Algoet et al., 2018; Grandjean et al., 2018; Vassiliadis et al., 2018; Wilhelm et al., 2016). Adding a short interval between both pulses allows to avoid direct electromagnetic interference between the two coils, while preventing transcallosal interactions to occur between motor areas. This double-coil_1ms_ technique has proved useful to yield MEPs in muscles of both hands on each trial that are comparable to those obtained using single-coil TMS, whether elicited at rest (Grandjean et al., 2018) or during movement preparation (Vassiliadis et al., 2018), and whatever the order of stimulation (left-right or right-left M1); here, the first pulse was systematically applied over right M1. Both pulses were delivered through small figure-of-eight coils (wing internal diameter 35 mm); the coil stimulating left M1 was connected to a Magstim Bistim2 magnetic stimulator (Magstim, Whitland, Dyffed, UK), while the one stimulating right M1 was plugged into a Magstim 200 magnetic stimulator. Critically, both stimulators delivered monophasic pulses.

The coils were placed tangentially on the scalp with the handle pointing backward and laterally at 45° angle away from the midline, approximatively perpendicular to the central sulcus. For each M1, the optimal coil position for eliciting MEPs in the contralateral first dorsal interosseous (FDI), an index finger abductor, was identified and marked on a head cap placed on the participant’s scalp to provide a reference mark throughout the experiment (Derosiere et al., 2017a, 2017b; Vandermeeren et al., 2002; Zenon et al., 2018). The resting motor threshold (rMT) was determined as the minimal TMS intensity required to evoke MEPs of 50 μV peak-to-peak in the relaxed FDI muscle in 5 out of 10 consecutive stimulations. Across participants, the rMTs corresponded to 57.1 ± 10.7 % and 39.8 ± 7.1 % of the maximum stimulator output for the left and right M1, respectively.

The rMT difference between the two M1 was expected given the use of different stimulators (Duque et al., 2012). For each hand, the intensity of TMS throughout the experiment was always set at 115 % of the individual rMT.

### 2.5. EMG recording

EMG activity was recorded from surface electrodes (Ambu Blue Sensor NF-50-K Neuroline, Medicotest, Oelstykke, Denmark) placed over the FDI muscle of the left and right hands, allowing to extract MEPs in both the responding and the non-responding homologous muscles on each trial. Note that for all participants, stimulating the hotspot for the FDI also elicited reliable MEPs in the abductor digiti minimi (ADM), a pinkie abductor muscle. These MEPs were also considered bilaterally in the present study and provided us with a measure of CSE in an irrelevant muscle.

EMG data were collected for 3200 ms on each trial, starting 200 ms before the TMS pulse. The raw EMG signals were amplified (gain, 1K) and digitized at 2000 Hz for off-line analysis. The latter consisted in extracting the peak-to-peak amplitude of MEPs recorded in the FDI and ADM muscles. In order to prevent contamination of MEP measurements by significant fluctuations in background EMG, trials with any EMG activity larger than 100 μV in the 200 ms window preceding the TMS pulse were excluded from the analysis (Derosiere et al., 2015). Trials in which participants had provided a wrong response were also removed from the MEP data set. The remaining MEPs were classified according to the experimental condition within which they were elicited; for each condition, we excluded trials with peak-to-peak MEP amplitudes exceeding 2.5 SD around the mean. After trimming the data for errors, background EMG activity and outliers, we only kept subjects in which a minimum of 8 MEPs remained to assess CSE in each condition (see below exclusion rate).

### 2.6. Data selection for MEPs elicited during the RT period

An important goal of the present experiment was to identify changes in CSE occurring during movement preparation in the simple and choice RT blocks, with the main objective to evaluate the influence of action selection requirements on preparatory inhibition. To do so, we focused on two phases of the RT period, considering changes occurring either early on, shortly after the onset of the imperative signal, or later on, close to movement onset.

To assess changes occurring at the later stage of the RT period, we pooled together all MEPs elicited by a TMS pulse applied 145 to 45 ms before movement initiation. On average, these MEPs, called MEP_LATE_, were elicited at a comparable time with respect to movement onset in the two block types (simple: – 87.84 ± 1.42 ms and choice: −85.30 ± 1.49 ms; t_953_ = −1.23; p = 0.22). To characterize changes occurring earlier on, we used the MEPs elicited at TMS_RT-PERIOD-1_ (i.e. elicited 80 ms after the imperative signal; called MEP_EARLY-1_) and those elicited at TMS_RT-PERIOD-2_ (i.e. elicited 130 ms after the imperative signal; MEP_EARLY-2_). Note that the MEP_EARLY-2_ were only considered for the choice RT blocks as they often fell in the category of the MEP_LATE_ for most trials in the simple blocks, given the shorter RTs in this context. Importantly, all trials in which an MEP_EARLY-1_ or MEP_EARLY-2_ could also be categorized as an MEP_LATE_ (because subjects responded particularly fast) were excluded from the MEP analyses. This resulted in the exclusion of a few subjects in whom the minimum number of trials (i.e. 8) was not reached in all conditions. As a consequence, the analysis of MEPs during the RT period of simple and choice RT blocks was run on 16 and 13 subjects, respectively (instead of 18 subjects). Note that all subjects excluded from this analysis were nevertheless included for the analysis of MEPs probed outside this period, i.e. at TMS_BASELINE-OUT_, TMS_BASELINE-IN_ and TMS_IMPERATIVE_.

### 2.7. Statistical analyses

#### Behavioral data

To characterize behavior in the rolling ball task, we focused on the RTs (participants made very few errors in the task). For this analysis, we only considered trials in which the TMS pulse was applied before the RT period and had thus a minor impact on behavior (Duque et al. 2014; Klein et al. 2015). A three-way repeated measure (RM) analysis of variance (ANOVA) was computed on these RTs, with BLOCK-TYPE (Simple, Choice), RESPONDING-HAND (Left, Right), and TMS-TIMING (TMS_BASELINE-IN_, TMS_IMPERATIVE_) as within-subject factors.

#### CSE data

First, we focused on FDI and ADM MEPs elicited at rest, when subjects were not preparing a response, considering MEPs obtained outside the blocks (TMS_BASELINE-OUT_) and those recorded within the blocks (TMS_BASELINE-IN_). For this analysis, MEPs were classified according to whether they were elicited (1) in a hand that was always responding in the related block : left (right) MEPs in the context of simple RT blocks requiring left (right) hand responses, (2) in a hand that could potentially be responding: left and right MEPs in the context of choice RT blocks, or (3) in a hand that was always non-responding: left (right) MEPs in the context of simple RT blocks requiring right (left) hand responses. The raw amplitude of FDI and ADM MEPs (mV) was then analyzed using two separate RM ANOVAs (one for each muscle), with MEP-SIDE (Left, Right), HAND-FUNCTION (Always responding, Potentially responding, Always non-responding), and TMS-TIMING (TMS_BASELINE-OUT_, TMS_BASELINE-IN_) as within-subject factors.

Second, we focused on FDI and ADM MEPs probed at TMS_IMPERATIVE_. Those MEPs were expressed in percentage of MEPs elicited at TMS_BASELINE-IN_ and were used to evaluate whether any MEP change occurred at the onset of the imperative signal, and, if any, whether this change would depend on the function of the hand in the task. To do so, two RM ANOVAs (one per muscle) were computed on these data, with MEP-SIDE (Left, Right) and HAND-FUNCTION (Always responding, Potentially responding, Always non-responding) as within-subject factors. Moreover, in order to assess the significance of changes in CSE in each sub-condition, one-sample t-tests comparing MEPs to a constant value of 100 (i.e. to TMSbaseline-in) were performed.

Third, to characterize the evolution of CSE during action preparation, four separate RM ANOVAs (one per block type and per muscle) were performed on MEPs (expressed in percentage of MEPs elicited at TMS_BASELINE-IN_) elicited during the RT period, using RESPONDING-SIDE (Left, Right), MEP-SIDE (Left, Right), and MEP-TIMING (MEP_EARLY-1_ and MEP_LATE_ in simple blocks; MEP_EARLY-1_, MEP_EARLY-2_ and MEP_LATE_ in choice blocks) as within-subject factors. Additionally, one-sample t-tests were used to compare MEPs in each sub-condition to a constant value of 100 (i.e. to TMS_BASELINE-IN_). Due to the exclusion of few subjects, those analyses were performed on 16 subjects for the simple RT blocks, and on 13 subjects for the choice RT blocks.

Finally, we performed a last analysis to make direct comparisons between simple and choice RT blocks. To do so, we focused on a condition showing some preparatory inhibition; that is, on MEP_LATE_ in a non-responding hand (see Result section). To analyze these data, we used a RMANOVA with MUSCLE (FDI, ADM), BLOCK-TYPE (Simple, Choice) and MEP-SIDE (Left, Right) as within-subject factors. This analysis was performed on the 11 subjects for whom we had the full set of data in both simple and choice RT blocks.

The Fisher’s Least Significant Difference (LSD) method was used to run post-hoc comparisons. All of the data are expressed as mean ± SE and the statistical significance was set at *p* < 0.05. Analyses were carried out using Statistica 10 (StatSoft, Cracow, Poland).

## Results

### Behavioral data

Overall, the mean RT in the simple blocks was 244 (SE = 8.6) and 237 ms (SE = 7.1) for responses performed with the left and right hand, respectively. The corresponding means for the choice blocks were 337 (SE = 8.1) and 322 ms (SE = 6.6). As evident from the numbers and as confirmed by the main effect of BLOCK-TYPE (F_1,17_= 199.53; *p* < 0.001), RTs were considerably shorter in the simple than in the choice blocks (Figure 2). Moreover, we observed a significant effect of RESPONDING-HAND (F_1,17_= 8.77*;p* < 0.05): participants were faster to respond with their right than with their left hand, regardless of the block type (BLOCK-TYPE x RESPONDING-HAND interaction; F_1,17_=1.26; p = 0.28).

**Figure 2.**
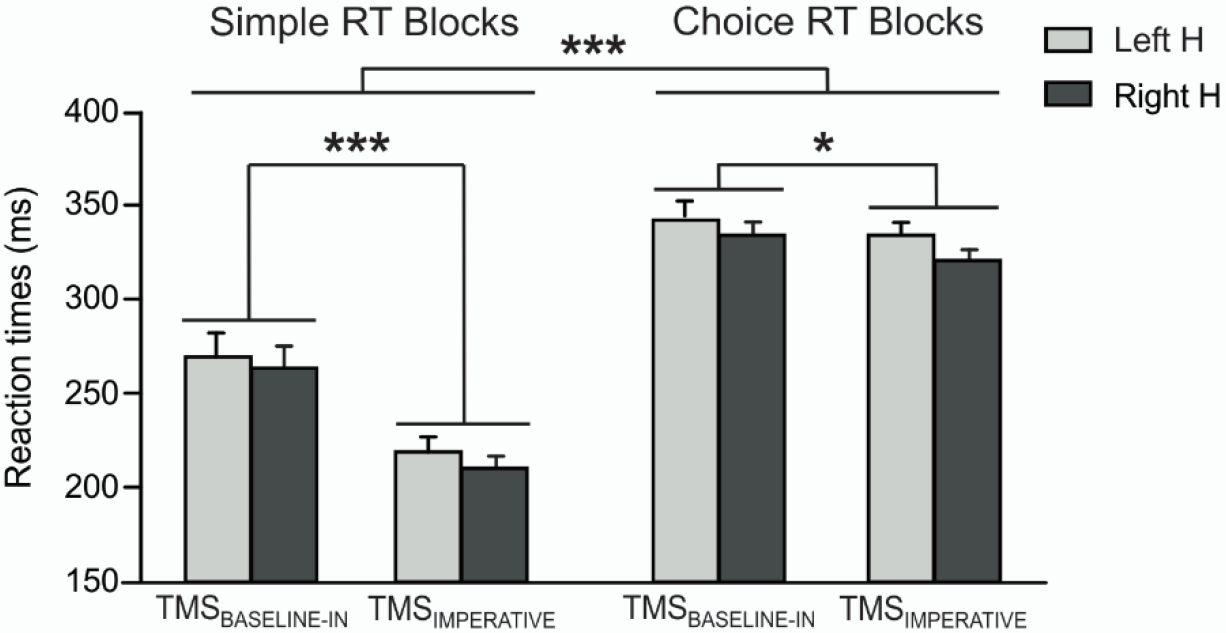
Reaction times (ms) for responses provided with the left (light grey) and right hands (H, dark grey) in the simple and choice blocks in trials in which the TMS pulses were either applied during the blank screen (i.e. TMS_BASELINE-IN_) or at the onset of the imperative signal (i.e. TMS_IMPERATIVE_). \**p* < 0.05 and \*\*\**p* < 0.001: significantly different.

RTs were also influenced by the TMS-TIMING (F_1,17_= 21.86; *p* < 0.001). As such, RTs were shorter when the TMS pulse was applied at the onset of the imperative signal (TMS_IMPERATIVE_) compared to when the pulse was applied during the blank screen (TMS_BASELINE-IN_). Hence, a pulse applied simultaneously with the imperative signal boosted the release of the future response.

Interestingly, this speed-up effect depended on the block type (BLOCK-TYPE x TMS-TIMING interaction; F_1,17_=10.72; p < 0.05) as well as on the hand used to respond (RESPONDING-HAND x TMS-TIMING interaction; F_1,17_=4.88; p < 0.05). As such, it was more prominent in simple than in choice blocks (p < 0.001 and p < 0.05, respectively) and stronger for right than left hand responses (p < 0.001 and p < 0.05, respectively). The BLOCK-TYPE x RESPONDING-HAND x TMS-TIMING interaction was not significant (F_1,17_=0.459; p = 0.51).

#### CSE data

#### MEPs elicited at TMSBASELINE

First, we considered MEPs elicited at rest, either outside or within the blocks (TMS_BASELINE-OUT_ and TMS_BASELINE-IN_; Figure 3A). Overall, the amplitude of FDI MEPs probed at TMS_BASELINE-OUT_ was 1.54 (SE = 0.24) and 1.83 mV (SE = 0.34) in the left and right hands, respectively. When elicited at TMS_BASELINE-IN_, FDI MEPs equaled 2.40 (SE = 0.43) and 2.78 mV (SE = 0.45) in the corresponding conditions. There was no main effect of MEP-SIDE (F_1,17_ = 0.90; p = 0.36). In contrast, as evident in the numbers and confirmed by the main effect of TMS-TIMING (F_1,17_ = 19.13; p < 0.001), FDI MEPs elicited at TMS_BASELINE-IN_ were larger relative to those obtained at TMS_BASELINE-OUT_ (see Figure 3A, left panel). Hence, FDI MEP amplitudes were higher in the context of the task, as shown in previous studies (Duque et al., 2016; Vassiliadis et al., 2018). Interestingly, the MEP increase at TMS_BASELINE-IN_ depended on the function of the hand within which they were elicited (HAND-FUNCTION x TMS-TIMING interaction; F_2,34_=14.72; p < 0.001). As such, while FDI MEPs were comparable in all conditions when elicited outside the blocks (all p > 0.37), their amplitude at TMS_BASELINE-IN_ seemed positively related to the level of involvement of the hand in the task. Indeed, the largest FDI MEPs were observed in a hand that was always used to respond (i.e. the responding hand in the simple blocks; *p* < 0.001 compared to both other conditions). Moreover, FDI MEPs were higher in a potentially responding hand (i.e. any hand in the choice blocks) than in an always non-responding hand in the simple blocks (*p* < 0.01). These results did not depend on the side of the MEP recording (MEP-SIDE x HAND-FUNCTION x TMS-TIMING interaction; F_2,34_ = 0.07; *p* = 0.93).

**Figure 3.**
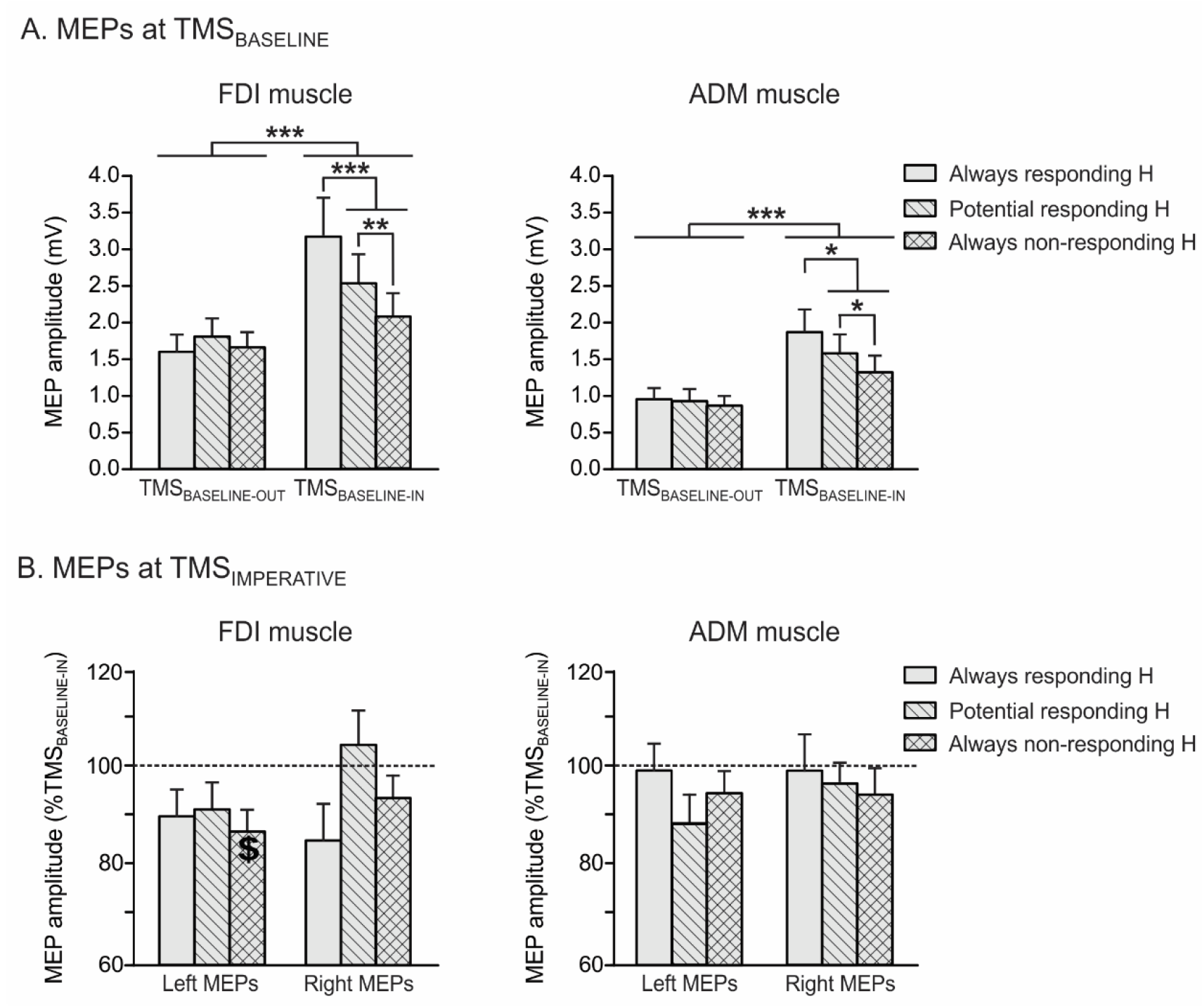
Amplitude of motor-evoked potentials (MEPs) recorded at rest (A) and at the onset of the imperative signal (B) in the first dorsal interosseous (FDI, left panel) and the abductor digiti minimi (ADM, right panel) muscles. MEPs recorded at rest were probed either during the blocks (TMS_BASELINE-IN_) or outside them (TMS_BASELINE-OUT_) and were expressed in mV, while MEPs recorded at the onset of the imperative signal were expressed in percentage of MEPs elicited at TMS_BASELINE-IN_. MEPs are shown for an always responding hand (i.e. the responding hand in the simple blocks; open bars), a potentially responding hand (i.e. both hands in the choice blocks; hatched bars), and an always non-responding hand (i.e. the non-responding hand in the simple blocks; crosshatched bars). Data are collapsed across both hands. \**p* < 0.05, \*\**p* < 0.01 and \*\*\**p* < 0.001: significantly different. $: significantly different from MEPs elicited at TMS_BASELINE-IN_. H = hand.

MEPs elicited in the ADM showed a similar pattern of changes as those reported above for the FDI. As shown on the right panel of Figure 3A, ADM MEPs were considerably greater when elicited at TMS_BASELINE-IN_ (1.59 ± 0.25 mV) than at TMS_BASELINE-OUT_ (0.91 ± 0.14 mV; p < 0.001). In addition, this increase in CSE also depended on the function of the hand within which they were recorded, although here the ADM was always irrelevant in the task (HAND-FUNCTION x TMS-TIMING interaction F_2,34_=4.36; p < 0.05).

#### MEPs elicited at TMS_IMPERATIVE_

Then, we focused on MEPs probed at the onset of the imperative signal (Figure 3B). These MEPs were expressed in percentage of MEPs elicited at TMS_BASELINE-IN_ and were considered in order to test for any change in CSE that could be due to the anticipation of the imperative signal. There was no main effect of the factor MEP-SIDE neither for the FDI (F_1,17_ = 1.90; *p* = 0.19) nor for the ADM (F_1,17_ = 0.36; *p* = 0.56). Moreover, we did not observe any effect of the factor HAND-FUNCTION (F_2,34_ = 1.18; *p* = 0.32 and F_2,34_ = 0.56; *p* = 0.58 for the FDI and ADM). Finally, MEPs at TMS_IMPERATIVE_ were comparable to those elicited at TMS_BASELINE-IN_ in all but one condition (see Figure 3B), indicating very little anticipation in this task.

#### MEPs elicited during the RT period

MEPs probed during the RT period are depicted on Figure 4 (FDI) and Figure 5 (ADM). These MEPs were expressed in percentage of MEPs elicited at TMS_BASELINE-IN_ for data analyses.

**Figure 4.**
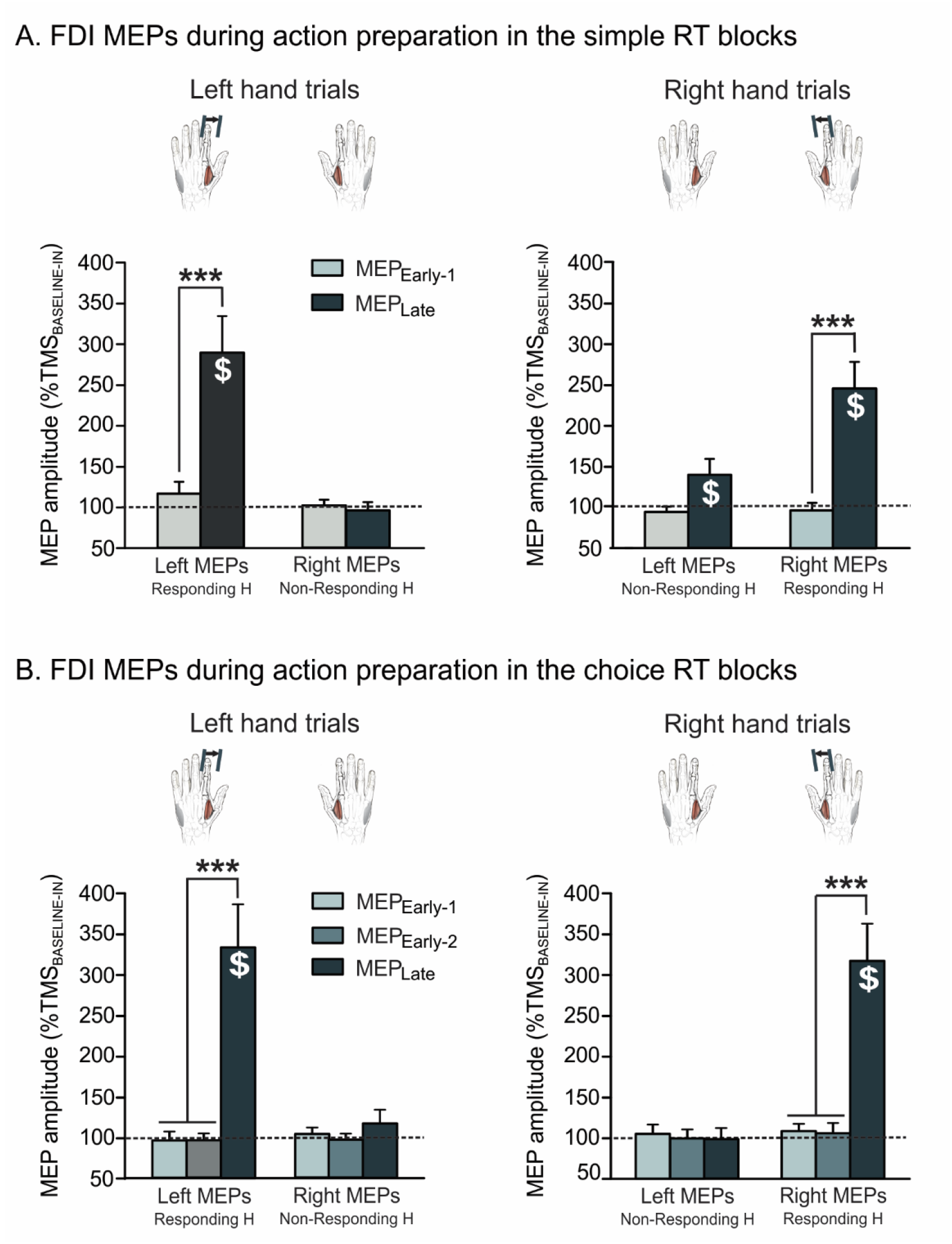
Amplitude of motor-evoked potentials (MEPs) elicited during the RT period (expressed in percentage of MEPs elicited at TMS_BASELINE-IN_) in the first dorsal interosseous (FDI; agonist muscle in the task) of the left and right hands. These normalized MEPs are shown for the simple (A) and choice (B) RT blocks and because they are recorded preceding both left responses (left panel) and right responses (right panel), they always either reflect changes occurring in a responding hand (e.g. Left MEPs in Left hand trials) or a non-responding hand (e.g. Left MEPs in right hand trials). \*\*\**p* < 0.001: significantly different. $: significantly different from MEPs elicited at TMS_BASELINE-IN_. H = hand. MEP_EARLY-1_ = MEPs elicited 80 ms after the onset of the imperative signal. MEP_EARLY-2_ = MEPs elicited 130 ms after the onset of the imperative trial. MEP_LATE_ = MEPs elicited 145 to 45 ms before movement onset.

**Figure 5.**
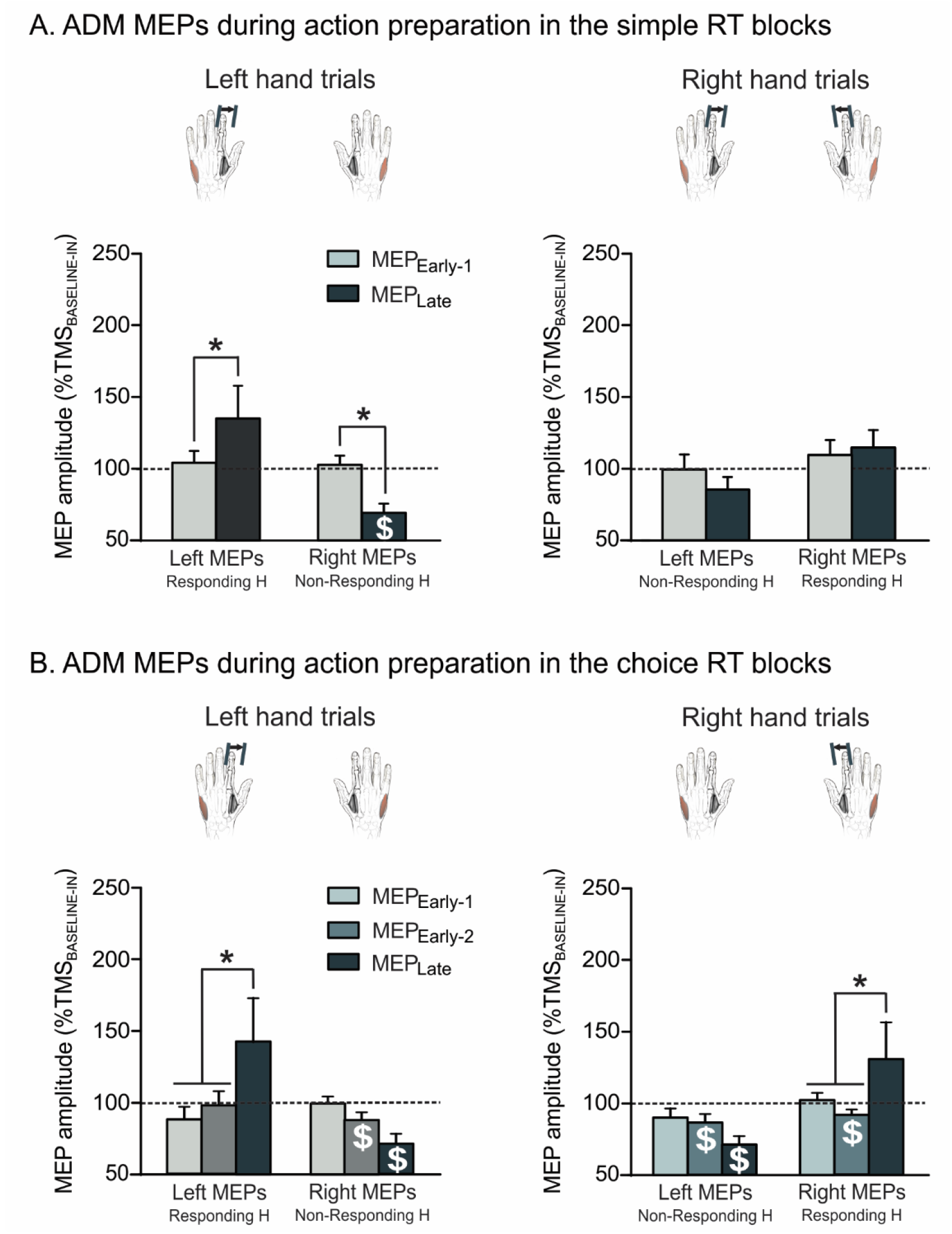
Amplitude of motor-evoked potentials (MEPs) elicited during the RT period (expressed in percentage of MEPs elicited at TMS_BASELINE-IN_) in the abductor digiti minimi (ADM; irrelevant muscle in the task) of the left and right hands. These normalized MEPs are shown for the simple (A) and choice (B) RT blocks and because they are recorded preceding both left responses (left panel) and right responses (right panel), they always either reflect changes occurring in a responding hand (e.g. Left MEPs in Left hand trials) or a non-responding hand (e.g. Left MEPs in right hand trials. \**p* < 0.05 and \*\**p* < 0.01: significantly different. $: significantly different from MEPs elicited at TMS_BASELINE-IN_. H = hand. MEP_EARLY-1_ = MEPs elicited 80 ms after the onset of the imperative signal. MEP_EARLY-2_ = MEPs elicited 130 ms after the onset of the imperative trial. MEP_LATE_ = MEPs elicited 145 to 45 ms before movement onset.

First, we analyzed MEPs obtained from the FDI muscle in simple RT blocks (Figure 4A). We observed a main effect of the factors MEP-TIMING (F_1,15_=24.94; p < 0.001) and MEP-SIDE (F_1,15_=4.59; p < 0.05), yet these effects depended on other factors. For instance, the effect of MEP-SIDE depended on the RESPONDING-SIDE (F _1,15_=_16_._86_; p < 0.001). This interaction was expected: as such, left hand MEPs were larger when the simple RT task required left than right hand responses, whereas right hand MEPs were larger preceding right than left hand responses. Hence, MEPs during the RT period were larger when they were obtained from the responding FDI, compared to when they occurred in the non-responding hand. Also predicted is the fact that this effect depended on the time at which MEPs occurred (RESPONDING-SIDE x MEP-SIDE x MEP-TIMING interaction; F_1,15_=24.67; p < 0.001); it was evident for MEP_LATE_, falling close to movement initiation (all p < 0.001 when comparing responding and non-responding conditions for left and right hand MEPs), but not at an earlier stage of the RT period, for MEP_early-1_ (all p > 0.45). In fact, the amplitude specifically increased from MEP_EARLY-1_ to MEP_LATE_ in the responding hand (all p < 0.001), while MEPs remained unchanged over time in the non-responding hand (all p > 0.13 when comparing MEP_EARLY-1_ and MEP_LATE_). Accordingly, additional analyses did not reveal any suppression of FDI MEPs in simple RT blocks (with respect to baseline), even when they were elicited in a non-responding hand. If anything, non-responding MEPs showed some facilitation close to movement onset, but this effect only occurred in right hand trials (Nonresponding left MEP_LATE_ t_15_=2.25, *p* < 0.05 when compared to a constant value of 100; all other p > 0.23).

Overall, the same dynamics of FDI MEP changes were observed in the choice RT blocks (Figure 4B), with the RM ANOVA revealing similar interactions, including the RESPONDING-SIDE x MEP-SIDE x MEP-TIMING interaction (F_2,24_=42.34; p < 0.001). FDI MEPs elicited during the RT period showed a specific increase when they were recorded from the responding hand (all p < 0.001) but not when they were obtained from the non-responding hand (all p > 0.46), resulting in larger MEPs close to movement initiation in the former than in the latter condition (all p < 0.001). Finally, here too, we did not observe any MEP suppression during the RT period (all p > 0.24). Yet, unlike simple blocks, we did not observe any facilitation of non-responding left MEP_LATE_.

Interestingly, the RESPONDING-SIDE x MEP-SIDE x MEP-TIMING interaction was also significant for MEPs recorded from the ADM (see Figure 5), i.e. a muscle that was always irrelevant in the task. This was true for both types of blocks (F_1,15_ = 13.47; p < 0.01 and F_2,24_=6.76; p < 0.01 for the simple and choice blocks, respectively). When probed in the responding hand, similar to the observations made on the FDI, ADM MEPs were larger close to movement initiation relative to earlier timings (all MEP_LATE_ p < 0.05, except when elicited in the right hand in the simple blocks: p = 0.66). This finding indicates that in most conditions, preparation of the required movement produced, to some extent, facilitation of irrelevant effectors in the responding hand. More importantly, when probed in the non-responding hand, ADM MEPs showed a significant decrease in their amplitude close to movement onset. As such, MEP_LATE_ were significantly suppressed (with respect to baseline) when probed in the right hand (both block types t < – 4.51; p <0.001) and in the left hand (only in choice blocks t_12_ = −4.46, p < 0.001; not significant in the simple blocks, with t_15_ = −1.70, p = 0.11), contrasting with the absence of MEP suppression for the non-responding FDI. Note that the ADM also displayed some suppression when MEPs were elicited in the responding hand, before their facilitation close to movement onset. Even if this was only true in one condition, i.e. when MEP_EARLY-2_ were elicited in the right ADM in choice RT blocks (t_12_ = −3.12, p < 0.01). This finding suggests that motor suppression is not necessarily restricted to muscles of the non-responding hand.

Finally, a last analysis focused on FDI and ADM MEP_LATE_ in the non-responding hand, i.e. the condition showing some MEP suppression. Our reasoning was that if preparatory inhibition assists action selection, these MEP_LATE_ may be smaller in the choice relative to the simple blocks. Surprisingly, neither the factor BLOCK-TYPE (F_1,10_=_0_._01_; p = 0.96), nor the MUSCLE x BLOCK-TYPE interaction (F_1,10_=0.0173; p = 0.95) were statistically significant. Hence, having to select between two actions does not appear to entail further inhibition of the inappropriate response in the non-responding hand. However, this interpretation is challenged by the significant MUSCLE x BLOCK-TYPE x MEP-SIDE interaction (F_1,10_=5.48; p < 0.05). As evident on the left side of Figure 6, while FDI MEPs were lower in the choice than in the simple blocks when elicited in the left nonresponding hand (p < 0.01), the reverse effect was found in the right non-responding hand, with larger MEPs in the choice compared to the simple RT blocks (p < 0.01). The same trend was observed for the ADM, although the comparison between both block types did not reach significance for that irrelevant muscle (p>0.25 for left and right MEPs, see right side of Figure 6).

**Figure 6.**
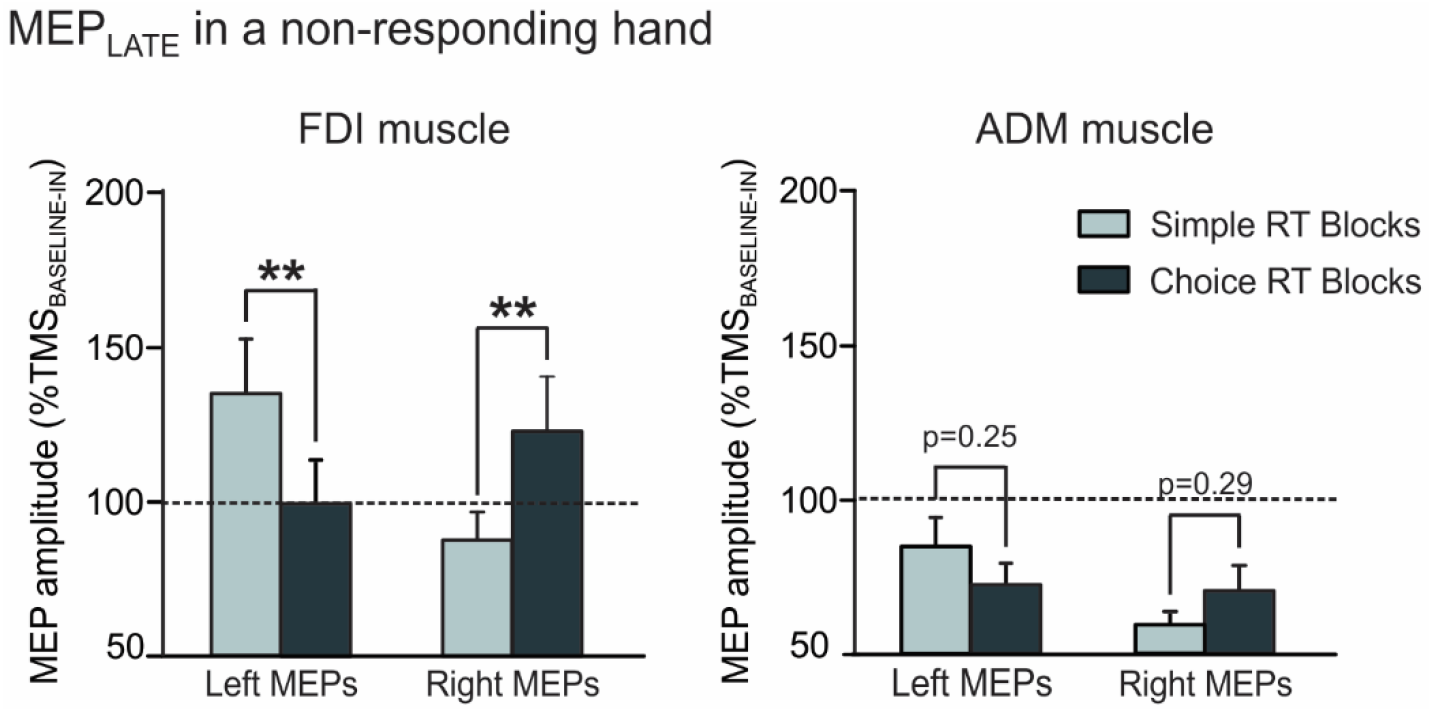
Amplitude of motor-evoked potentials (MEPs) elicited at a late stage of the RT period (called MEP_LATE_, falling 145-45 ms before movement onset; expressed in percentage of MEPs elicited at TMS_BASELINE-IN_) in the first dorsal interosseous (FDI, agonist muscle in the task; left panel) and abductor digiti minimi (ADM, irrelevant muscle in the task; right panel) of the nonresponding hand (i.e. Left MEPs in Right hand trials and Right MEPs in Left hand trials). These normalized MEPs are shown for the simple (A) and choice (B) RT blocks \*\**p* < 0.01: significantly different.

## Discussion

Recently, the prevalent idea that preparatory inhibition assists action selection has been challenged (Duque et al., 2017; Greenhouse et al., 2015; Hannah et al., 2018). Here, we aimed at addressing the contribution of preparatory inhibition to action selection by contrasting MEPs elicited during action preparation in a simple and a choice RT task within the same session. Critically, all other aspects of the task were identical in both settings and the experimental design minimized the influence of inhibitory processes such as proactive inhibition (no catch trials) or impulse control (no pre-cue, random intertrial interval).

The first result of interest here pertains to the mere presence of some MEP suppression in the nonresponding hand in the current setting. This aspect is crucial, as most previous TMS works reported suppression of the motor system using tasks in which other forms of inhibition might have operated. For instance, past experiments have often used task designs where the imperative signal is preceded by an alerting pre-cue (Duque et al., 2005; Leocani et al., 2000; Tandonnet et al., 2011). In such studies, MEPs are already drastically suppressed during the pre-imperative period, an effect that would allow them to withhold their response until the appropriate time, and could then persist in the non-responding effector. Moreover, past experiments have often included catch trials requiring subjects to anticipate the possibility of a “no-go” instead of the imperative signal. Although these trials are often rare (~5% of total number of trials, but see Greenhouse et al., 2015; Quoilin and Derosiere, 2015), they might have led to some proactive inhibition, contributing to the observed MEP suppression during action preparation in these studies. Here, by making the onset of the imperative signal less predictable and by avoiding catch trials, we minimized these other inhibitory sources, a claim further supported by the overall lack of MEP changes at the onset of the imperative signal. Hence, the MEP suppression here cannot be accounted for by such anticipatory or proactive inhibitory forms.

Intriguingly, MEP suppression only concerned the ADM, which was irrelevant in the task. Although the lack of suppression in the FDI might seem surprising, one has to keep in mind that MEPs represent global readouts of the excitability of the motor system, and that their amplitude results from multiple excitatory and inhibitory influences acting at the same time (Bestmann & Duque, 2016). Hence, it is possible that here, excitatory inputs prevailed for the FDI, resulting in a net facilitation of MEPs elicited in this muscle, although inhibitory influences may still have operated. Consistently, action preparation is known to entail a phenomenon referred to as crossed corticospinal facilitation (Carson et al., 2008), whereby the facilitation of the agonist effector (i.e. the responding FDI) spreads onto the homologous muscle in the resting hand (i.e. the nonresponding FDI). Yet, although this explanation is plausible, the lack of MEP suppression in the FDI is still in contradiction with previous work from our group, in which we used a very similar task where the ability to predict the onset of the imperative was also greatly reduced (Duque et al., 2014). However, the pattern of CSE changes in the responding effector was also considerably different in that previous study. That is, while MEPs probed close to movement initiation were around three times larger than those probed at baseline in the current experiment, this increase was marginal (i.e. around 125 %) in the other one (Duque et al., 2014). This inconsistency may be due to differences in the biomechanical properties of the required movement in both studies, as CSE is known to correlate with the magnitude of the upcoming muscular contraction (Cos et al., 2014; MacKinnon and Rothwell, 2000). Accordingly, participants were required to perform their response “in the air” in our previous work (Duque et al., 2014), whereas they had to reach the inner metal edge of the response device in the present experiment. Hence, the vigor of the upcoming movement was probably greater in the latter case, leading to an enhanced facilitation in the agonist muscle. The crossed corticospinal facilitation may thus have been particularly strong here, preventing MEPs from exhibiting any inhibitory change in the non-responding homologous FDI muscle.

Interestingly, inhibition in the non-responding ADM was already evident in the simple version of the task, but only for MEPs probed in the right hand, i.e. in a condition in which participants had to systematically perform their response with the left index finger. At first sight, one might argue that the presence of MEP suppression in the simple RT task is against the action selection hypothesis. Yet, because all our subjects were right-handed, right index finger movements may be considered as prepotent, such as suggested by the significantly faster responses performed with the right than with the left hand. As a result, although there was no explicit selection requirement in the simple RT task, preparing responses with the left (non-dominant) hand might have still required some inhibition to suppress the prepotent response of the right (dominant) hand. This hypothesis is in line with the observation that non-responding MEPs were smaller preceding left than right hand responses in the simple RT task, both in the FDIs and the ADMs, a finding which is also consistent with prior studies (Klein et al., 2016; Poole et al., 2018; but see Leocani et al., 2000).

By contrast, when participants were explicitly required to select an action (i.e. in the choice RT task), motor inhibition was always observed in the non-responding hand, regardless of whether the response was provided with the left or the right hand. This effect can be explained following the same line as for the simple RT task. That is, here, prepotency concerns the two hands given that they were both directly in competition and equally probable. Hence, they both had to be silenced in case the other one was selected. In other words, we propose that preparatory inhibition of the non-responding hand reflects a process that assists action selection by suppressing motor activity of prepotent/competing effectors, resulting in MEP suppression in the choice RT task but also preceding non-dominant hand responses in the simple RT task.

Another notable finding of the current work was the presence of some MEP suppression in the responding hand, at an early phase of action preparation. Although this effect was only evident for the right ADM during the choice version of the RT task, it is consistent with the idea that inhibition for action selection is a global process, suppressing broadly the motor system as of the beginning of action preparation, before persisting in the non-responding muscles (Duque et al., 2017). As such, even if MEPs elicited in the responding effector increase during the premovement period, their amplitude is often found to be initially reduced (Duque et al., 2014; Klein et al., 2016). This early decrease is also present in the non-responding hand, but there, MEPs display then a further drop in amplitude (Duque et al., 2014; Leocani et al., 2000; Tandonnet et al., 2011). Finally, the fact that MEPs were found suppressed in the ADM, irrelevant in the task, is also consistent with a broad mechanism targeting several muscles irrespective of their function in the forthcoming response (Duque et al., 2014; Greenhouse et al., 2015; Quoilin et al., 2016).

Lastly, MEPs probed at rest during the task were larger compared to MEPs probed outside the blocks, consistent with previous reports (Duque et al., 2016; Vassiliadais et al., 2018). Interestingly, this increase in MEP amplitudes was proportional to the degree of involvement of the hand in the task. That is, the increase was strongest for MEPs elicited in an always responding hand, and was more pronounced for MEPs in a potential responding hand relative to an always non-responding hand. Hence, it is likely that potential respondants, or “affordances”, remained somewhat constantly preactivated in the task to help respond as fast as possible (Cisek 2007; Gibson 1979; Wickens et al., 1994). In particular, the strong preactivation of MEPs in the simple RT task (always responding hand) is likely to contribute to the rapidity of responses in this setting and to the fact that TMS_IMPERATIVE_ had such a strong boosting effect in this condition: RTs were 52 ms faster in trials involving a pulse at the onset of the imperative signal compared to trials in which the pulse fell at TMS_BASELINE-IN_.

In conclusion, by comparing MEP changes in a simple and a choice RT task, the present paper brings some arguments in favor of the action selection hypothesis. That is, our findings suggest that preparatory inhibition is generated whenever prepotent/competing effectors need to remain silent. Critically, this inhibition does not appear to be selectively directed at the non-responding prime-mover (to be suppressed). Rather, it seems to be global as MEP suppression was most evident when probed in an irrelevant muscle of the non-responding hand and on some occasion, it also concerned the responding hand. Our data also indicate that the use of a task requiring quite vigorous movements can prevent one from detecting inhibitory changes because excitatory influences directed at the prime-mover spill over onto non-responding muscles. Future experiments using techniques allowing to target specifically inhibitory circuits, such as paired-pulse TMS, should allow to consolidate the outcomes of the present study.

## Acknowledgements

This work was supported by grants from the “Fonds Spéciaux de Recherche” (FSR) of the Université Catholique de Louvain, the Belgian National Funds for Scientific Research (FRS-FNRS: MIS F.4512.14) and the “Fondation Médicale Reine Elisabeth” (FMRE). CQ was a postdoctoral fellow supported by the FNRS.

